# Sub-strain-Dependent Differences in Gut Barrier Permeability, Bacterial Translocation and Post-Stroke Inflammation in Wistar Rats

**DOI:** 10.1101/2025.11.13.688185

**Authors:** Cristina Granados-Martinez, Nuria Alfageme-Lopez, Manuel Navarro-Oviedo, Victor Mora-Cuadrado, David Sevillano-Fernandez, Luis Alou-Cervera, M. Encarnación Fernandez-Valle, Olivia Hurtado, Maria A. Moro, Ignacio Lizasoain, Jesus M. Pradillo

## Abstract

**Background:** Stroke induces profound neuroinflammation and systemic immune dysregulation, including disturbances in gut homeostasis. Experimental evidence suggests that intestinal barrier permeability (IBP) and bacterial translocation (BT) critically influence stroke outcomes. However, biological variability among commonly used rodent sub-strains has received limited attention.

**Methods:** In this pilot study, we compared post-stroke immune responses in two Wistar rat sub-strains obtained from different suppliers: RccHan (Envigo) and RjHan (Janvier). Following transient middle cerebral artery occlusion, animals were assessed 72 hours later and stratified according to the presence or absence of BT. Immune cell populations in blood and bone marrow were analyzed by flow cytometry, and leukocyte infiltration into ischemic brain tissue was quantified by immunohistochemistry.

**Results:** Both sub-strains developed significant infarcts and neurological deficits. RccHan rats displayed larger infarct volumes and more extensive BT across multiple organs. In contrast, RjHan rats exhibited BT mainly confined to mesenteric lymph nodes but showed greater IBP. Although dissemination was broader in RccHan rats, overall bacterial burden was slightly lower compared with RjHan, and extra-intestinal bacterial composition differed between groups. Notably, RjHan rats presented stronger systemic and central immune activation, with marked alterations in lymphocyte and monocyte populations and enhanced granulocyte and T cell infiltration within ischemic lesions.

**Conclusions:** These findings demonstrate that sub-strain origin profoundly influences post-stroke intestinal barrier integrity, bacterial dissemination, and immune responses. Considering sub-strain-related variability is essential to improve reproducibility and translational relevance in preclinical stroke research.

## Introduction

Ischemic stroke (IS) is one of the leading causes of morbidity and mortality worldwide and represents a major public health challenge [1]. Beyond the initial brain injury induced by arterial occlusion, secondary processes including excitotoxicity, oxidative stress, apoptosis, and inflammation contribute significantly to the progression of damage [2,3]. The post-stroke inflammatory response is not confined to the central nervous system: systemic immune activation, secondary immunosuppression, and barrier dysfunction are now recognized as decisive factors in clinical outcome [4]. Within this context, the gut–brain axis has gained increasing attention, as the intestine emerges as a particularly vulnerable target organ after acute brain injury. Cerebral ischemia induces alterations in gastrointestinal motility, microbiota composition, and intestinal barrier permeability (IBP) [5,6]. The disruption of epithelial tight junctions facilitates bacterial translocation (BT) and the passage of microbial products into the bloodstream and peripheral organs such as the spleen and liver [7,8]. This phenomenon not only increases the risk of post-stroke infections but also amplifies the systemic inflammatory response and may exacerbate secondary brain damage [9,10]. Experimental models have shown that modulation of the microbiota or preservation of intestinal integrity significantly influences infarct size, leukocyte infiltration, and neurological recovery [11–13]. Despite the importance of these findings, potential sources of variability affecting the gut–brain axis in stroke remain insufficiently explored. The Wistar rat is one of the most widely used strains in preclinical stroke research, particularly in the transient middle cerebral artery occlusion (MCAO) model, owing to its availability, docility, and reproducibility [14]. However, physiological responses and even ischemic outcomes may differ between rat strains or sub-strains [15–17]. Moreover, several studies have demonstrated that animals with the same genetic designation but obtained from different commercial vendors may exhibit divergences in immune phenotypes, behavior, and gut microbiota composition [18–20]. These “vendor effects” represent a source of experimental variability that can compromise reproducibility and translational validity [21]. To date, however, it remains unclear how the origin of Wistar rats from different breeders might influence key stroke-related processes such as IBP, BT and systemic and cerebral inflammation. Janvier Labs and Envigo (currently Inotiv) maintain independent colonies of Wistar rats with distinct breeding histories. While the original strain was established at the Wistar Institute in 1906, the long-term separation of breeding nuclei and vendor-specific husbandry practices have resulted in accumulated genetic and microbiological divergence [22,23]. Consequently, although both animals are designated as “Wistar,” they are not genetically identical and may display differences in critical biological parameters. The present work was designed to explore potential differences in IBP, BT, inflammatory responses, and brain injury between Wistar rats from Janvier and Envigo. By identifying such differences, our aim is to select the most appropriate model for future studies, thereby improving reproducibility and the translational relevance of preclinical stroke research [24].

## Methods

Details of materials and procedures are available in online Supplemental Material. Data supporting the findings are available from corresponding authors upon reasonable request.

### Animals

Wistar rats were obtained from two commercial suppliers: Janvier Labs (Wistar stock maintained by Janvier; RjHan:WIS) and Inotiv/Envigo (RccHan:WIST, commonly referred to as “Wistar Han,” maintained from the original Hannover nucleus). All experiments were conducted in male Wistar rats weighing 250–300 g and aged 8–12 weeks. To minimize potential vendor-related biases, animals were acclimated in our facility for at least 14 days prior to the start of the studies [24], maintained under identical housing conditions (light/dark cycle, diet, bedding type, and environmental enrichment), and the order of interventions was randomized across batches from each supplier.

All procedures complied with the European Communities Council Directive (86/609/EEC), were approved by the Ethics Committee on Animal Welfare of the Complutense University (PROEX No. 305/19 and 219.3/24) and are reported in accordance with the ARRIVE guidelines. Efforts were made to reduce the number of animals using prior experience and statistical tools (http://www.biomath.info), and no mortality was observed in any of the animals.

### Experimental groups, focal cerebral ischemia and neurological evaluation

All experiments were conducted in a randomized (coin toss) and blinded manner. The experimental groups included naïve rats and rats subjected to middle cerebral artery occlusion (MCAO). Based on unpublished results from our laboratory using this experimental stroke model, together with previous evidence indicating that the BT process occurs between 48–72 hours after stroke [7,8], all animals in this study were sacrificed, and all determinations were performed, at 72 h after experimental IS. Following the analysis of bacterial growth in peripheral organs, each animal was further classified as either exhibiting bacterial translocation (BT) or no bacterial translocation (NBT).

Focal cerebral ischemia was induced by tandem permanent occlusion of the left common carotid artery (ligation) and the left MCA (electrocoagulation). Neurological function was assessed both in naïve animals and, in MCAO rats, at 72 h post-surgery using standardized motor and behavioral scales (see Supplemental Material for details).

### Infarct size and GBD determination by MRI

Infarct volume was assessed 72h post-IS by T2-weighted MRI (Bruker 1T). IBP was evaluated 72h post-IS in naïve and untreated MCAO rats using T1-weighted abdominal MRI with oral contrast agents (CAs; mannitol and MnCl_2_·4H_2_O). Rats were fasted, anesthetized, and monitored during imaging on a Bruker 4.7T system. Intestinal damage was quantified via contrast leakage using ImageJ. Full MRI details are in Supplemental Material.

### Bacterial translocation analysis

BT was assessed by microbiological cultures from mesenteric lymph nodes (MLN), liver, spleen, and lungs 72h post-surgery. Tissues were aseptically collected, homogenized, plated on selective media under aerobic/anaerobic conditions. Colony-forming units (CFUs) were counted and reported as log_10_ CFU/g. Bacterial classification followed the International Code of Nomenclature of Prokaryotes (ICNP). Further details in Supplemental Material.

### Determination of peripheral and central inflammation

Peripheral inflammation was evaluated by flow cytometry analysis of immune cell populations in blood and bone marrow (BM) from naïve animals and stroke animals at 72 h. Cells were stained with specific antibody panels and analyzed using FlowJo software. Central inflammation was assessed by immunofluorescence on brain sections from naïve and 72 h stroke animals, stained for granulocytes, T cells (CD3), and endothelial markers (RECA-1). Images were acquired using confocal microscopy and quantified with ImageJ software. Additional methodological details are provided in the Supplemental Material.

### Statistical analysis

Data are expressed as mean ± standard deviation (SD) for variables with normal distribution, or as median and interquartile range (IQR) for non-normally distributed variables. The normality of the data was assessed using the Kolmogorov-Smirnov test. For comparisons between two independent groups, Student’s t-test was applied for parametric data and Mann-Whitney for non-parametric data. When more than two groups were analyzed, for non-parametric data Krustal-Wallis test was applied followed by Dunn’s post hoc multiple comparisons test. Shannon and Simpson indices were used to evaluate TB diversity across organs. The Shannon index quantified the overall dispersion of TB events among organs, whereas the Simpson index assessed the dominance.

All statistical analyses were conducted in GraphPad Prism 8.0.1 software and R (v4.4.2), and a P-value of <0.05 was considered statistically significant.

## Results

### Strain-dependent differences in infarct volume and BT following IS

IS resulted in significant infarct volumes in both sub-strains of animals, with RccHan rats developing larger lesions than RjHan rats (Figure 1A). At 72h post-stroke, both sub-strains showed pronounced motor and behavioral impairments compared with naïve controls; however, despite the greater infarct size in RccHan rats, the severity of functional deficits did not differ significantly between the two strains (Figure 1B).

**Figure 1.**
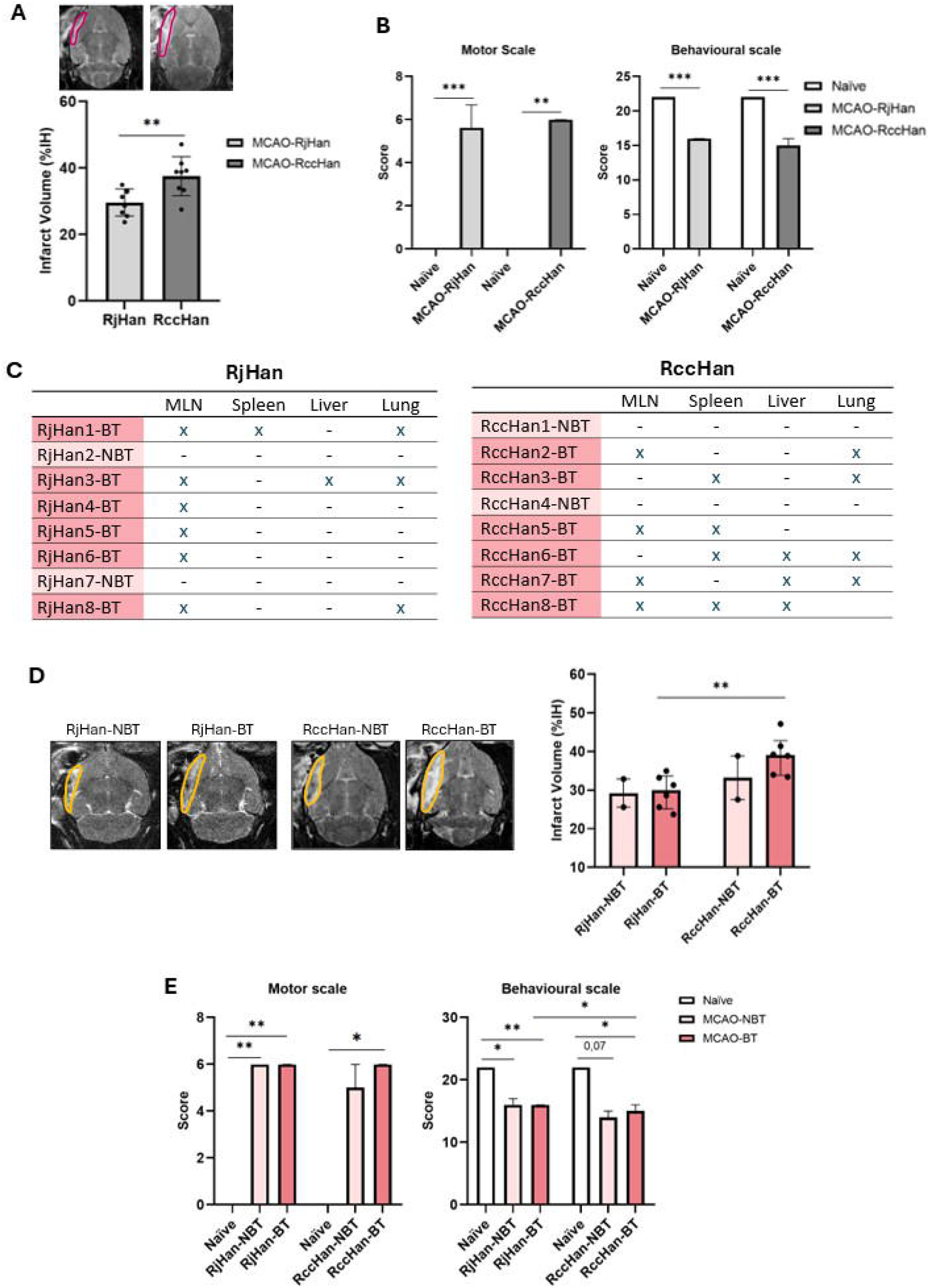
Infarct volume, stroke outcome, and bacterial translocation after experimental ischemic stroke in Wistar rats from two different commercial vendors. **A:** Infarct volume at 72 h post-surgery in MCAO-RjHan and MCAO-RccHan (n=8) animals. Infarct volume is expressed as the percentage of the ipsilateral hemisphere (%IH). Results are shown as mean ± SD; Student’s t-test, p < 0.01. **B:** Motor and behavioral characterization in naïve (n=4) and at 72 h in MCAO-RjHan and MCAO-RccHan (n=8) animals. Results are expressed as median (IQR). Statistical analysis was performed using the Kruskal–Wallis test followed by Dunn’s post hoc multiple comparisons test; ^**^p < 0.01, ^***^p < 0.001. **C:** Presence (x) / absence (-) of bacterial growth in different organs analyzed in each animal from the different experimental groups (*naïve*and MCAO). **D:** Infarct volume at 72 h post-surgery in RjHan-NBT (n=2), RjHan-BT (n=6), RccHan-NBT (n=2), and RccHan-BT (n=6) animals. Infarct volume is expressed as the percentage of the ipsilateral hemisphere (%IH). Results are shown as median (IQR); Kruskal-Wallis test followed by Dunn’s post hoc multiple comparisons test, ^**^p < 0.01. **E:** Motor and behavioral characterization in *naïve*(n=4) and at 72 h in RjHan-NBT (n=2), RjHan-BT (n=6), RccHan-NBT (n=2), and RccHan-BT (n=6) animals. Results are expressed as median (IQR). Statistical analysis was performed using the Kruskal–Wallis test followed by Dunn’s post hoc multiple comparisons test; ^*^p < 0.05, ^**^p < 0.01.

When BT was analyzed, our results showed bacterial growth in sterile organs (MLN, spleen, liver and lung) only in MCAO animals, occurring more frequently and with broader organ dissemination in RccHan than in RjHan rats (Figure 1C).

When infarct volume was analyzed according to translocation status, no significant differences were observed between both sub-strains of rats with or without BT. In contrast, direct comparison of the translocation-positive groups revealed that RccHan rats exhibited a wider bacterial dissemination than RjHam counterparts, consistent with their significantly larger infarct volumes (Figure 1D).

With respect to neurological outcome, both motor and behavioral scores were significantly impaired in MCAO animals of both sub-strains compared with *naïve*controls. However, BT did not further exacerbate functional deficits within either strain (Figure 1E).

### Bacterial Dissemination and Intestinal Barrier Integrity

In ischemic animals that developed bacterial translocation (BT), no statistically significant differences were detected between sub-strains overall; nevertheless, distinct trends were observed. While some RccHan rats exhibited bacterial presence in the mesenteric lymph nodes (MLN), the proportion of affected animals was consistently higher in the RjHan group. In contrast, bacterial growth in the spleen and liver was more frequently detected in RccHan rats, with no differences between sub-strains in the lung (Fig. 2A).

**Figure 2.**
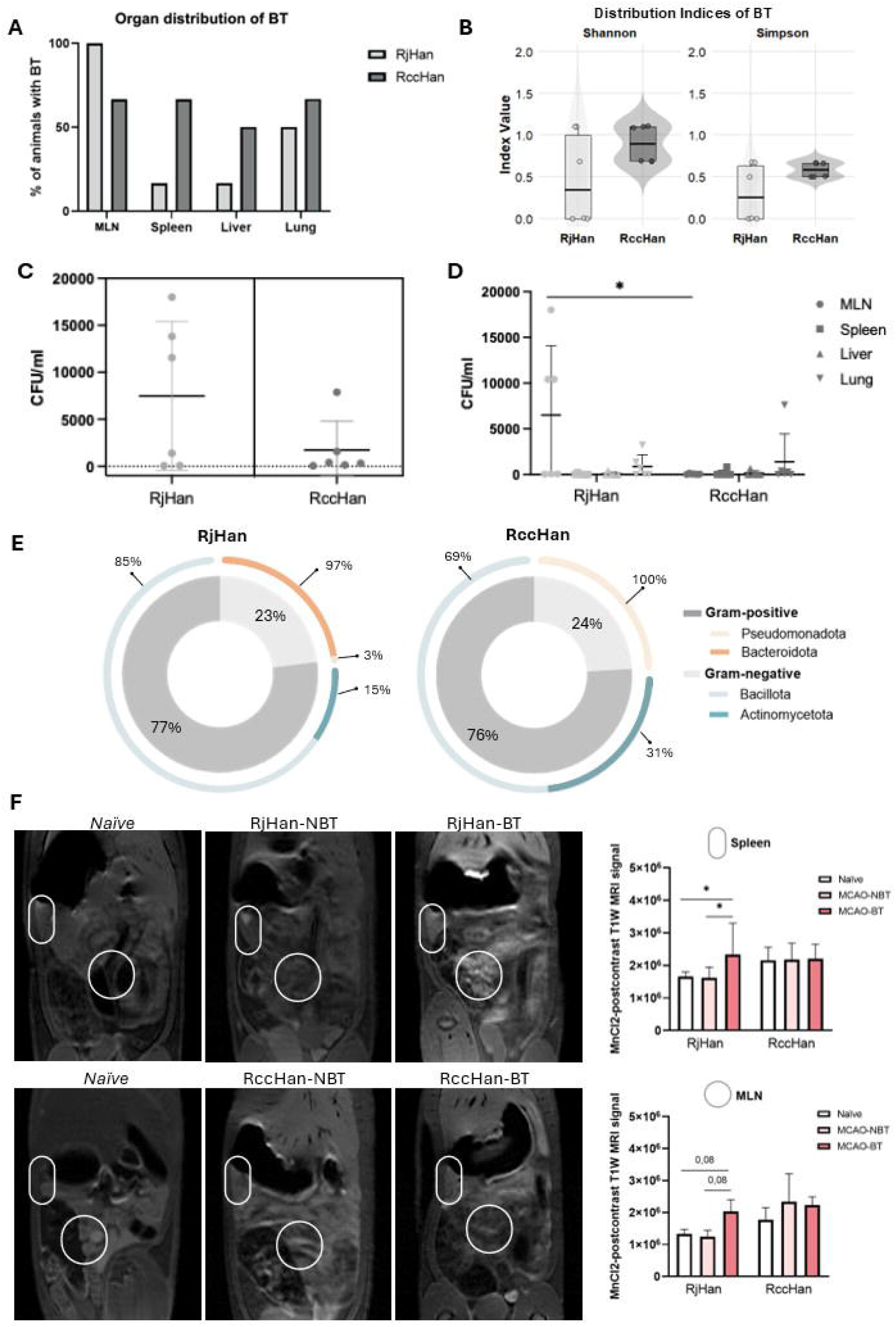
Effect of experimental ischemic stroke on bacterial translocation and intestinal barrier damage in animals from two different commercial vendors. **A:** Percentage of animals developing bacterial translocation in the different analyzed organs. Mann-Whitney Test. **B:** Diversity indices of BT distribution across organs (Shannon and Simpson indices) between both Wistar sub-strains. **C:** Representation of total bacterial load outside the intestine expressed as CFU/ml per animal in the RjHan-BT and RccHan-BT groups (n=6). **D:** Representation of bacterial load per analyzed organ expressed as CFU/ml per animal in the RjHan-BT and RccHan-BT groups (n=6). Results are shown as mean±SD; Mann-Whitney test, ^*^p<0.05. **E:** Relative abundance of total translocated Gram-positive and Gram-negative bacteria and bacterial phyla isolated from microbiological cultures of organs at 72 h after experimental IS in RjHan and RccHan (n=6). **F:** Intestinal permeability determined by measuring MnCl_2_-postcontrast T1W-MRI signal in spleen (oval) and MLN (circle) in naïve (n=4) and at 72 h after ischemic stroke in RjHan-NBT (n=2), RjHan-BT (n=6), RccHan-NBT (n=2), and RccHan-BT (n=6) animals. Results are shown as median (IQR); Kruskal–Wallis test followed by Dunn’s post hoc multiple comparisons test, ^*^p < 0.05.

Quantification of total CFU/ml across all sampled organs showed a trend toward lower bacterial burden in RccHan compared with RjHan rats, although this difference did not reach statistical significance (Fig. 2C). By contrast, organ-specific analysis revealed a significantly higher bacterial load in the MLN of RjHan rats relative to RccHan, confirming that BT was more pronounced in this organ and consistent with the more localized dissemination pattern observed in RjHan animals (Fig. 2D).

Taxonomic profiling of bacteria disseminated beyond the intestine showed that both sub-strains harbored phyla typically associated with the intestinal microbiota, and no differences were detected in the overall proportion of Gram-positive versus Gram-negative species. However, marked differences emerged at the phylum level. In RjHan rats, *Bacteroidota* predominated, followed by *Pseudomonadota*, whereas in RccHan rats *Bacteroidota* was absent and *Pseudomonadota* was the dominant phylum. Among Gram-negative taxa, *Bacillota* was the most abundant in RjHan rats, while RccHan rats exhibited a pronounced reduction in this phylum and, in particular, in *Actinomycetota* (Fig. 2E).

Finally, MRI analysis of IBP demonstrated significant barrier disruption in the spleen and a strong trend in the MLN of MCAO-RjHan rats with BT, findings that were absent in RccHan rats (Fig. 2F). This underscores the greater susceptibility of RjHan rats to BT-associated intestinal barrier impairment following ischemia.

### Peripheral and central inflammatory responses at 72 h after stroke

At 72h post-ischemia, both peripheral and central immune responses differed markedly between the two rat sub-strains and were further modulated by the presence of BT.

In the periphery, RjHan rats displayed stronger alterations than RccHan. In the BM, changes were generally more modest, although RjHan-BT animals exhibited clearer reductions in monocytes and increases in CD4^+^/CD8^+^ T cells compared with RccHan, suggesting enhanced mobilization or redistribution of immune populations (Fig. 3A). In peripheral blood, RjHan-BT rats showed significant increases in circulating CD4^+^ and CD8^+^ T cells together with a pronounced reduction in B cells and monocytes compared with *naïve* and non-BT groups, whereas these shifts were less evident or absent in RccHan (Fig. 3A, B).

**Figure 3.**
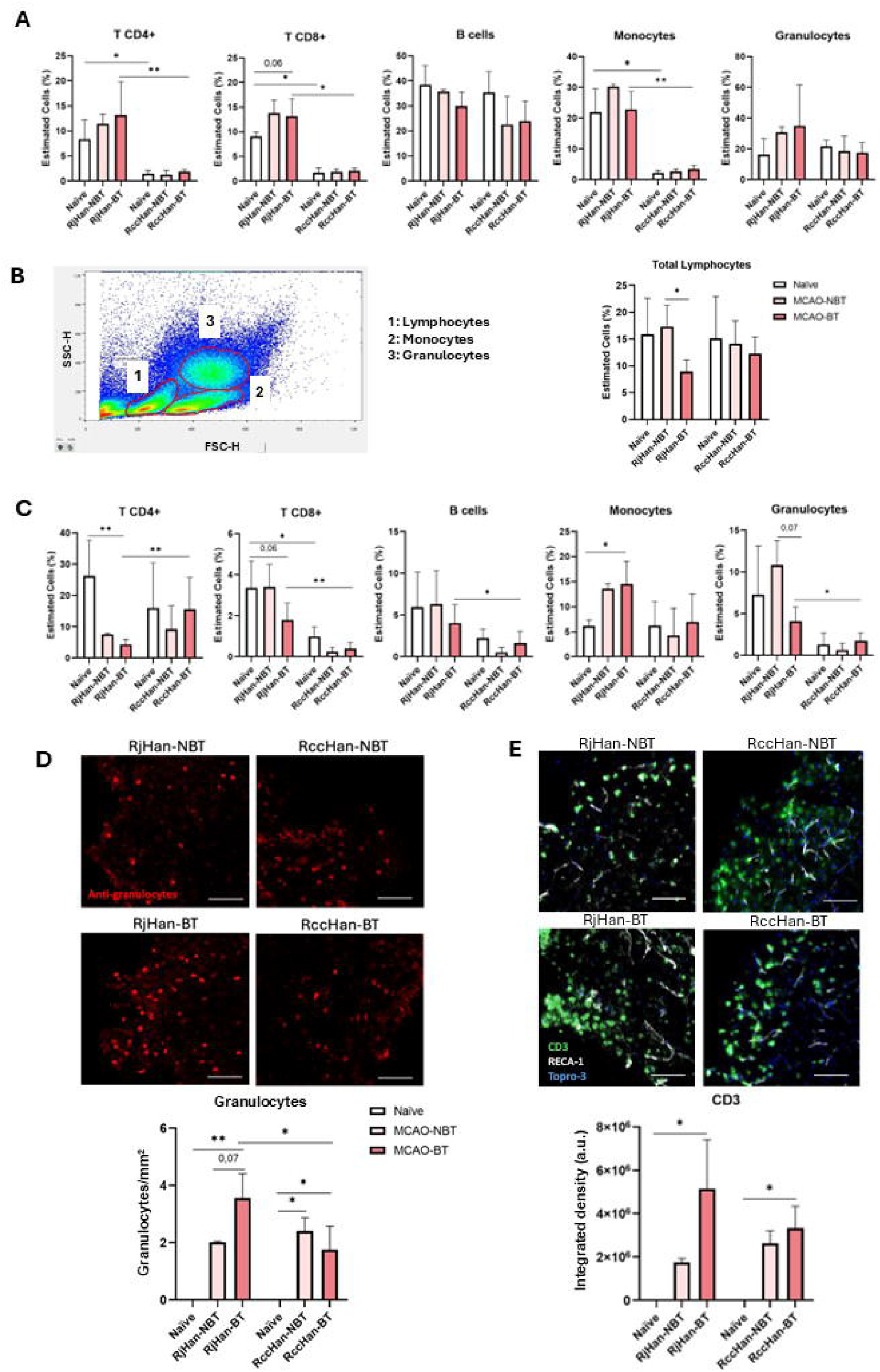
Impact of bacterial translocation on the peripheral and central inflammatory response at 72 h after experimental ischemic stroke. Immune cell levels assessed by flow cytometry in (**A**) bone marrow and (**B, C**) peripheral blood in *naïve* (n=4) and at 72 h post-surgery in RjHan-NBT (n=2), RjHan-BT (n=6), RccHan-NBT (n=2), and RccHan-BT (n=6) animals. Results are shown as median (IQR); Kruskal–Wallis test followed by Dunn’s post hoc multiple comparisons test, ^*^p < 0.05, ^**^p < 0.01. **D:** Quantification of granulocyte infiltration (granulocytes per mm^2^) in the ischemic brain at 72 h post-surgery in naïve (n=4) and at 72 h after ischemic stroke in RjHan-NBT (n=2), RjHan-BT (n=6), RccHan-NBT (n=2), and RccHan-BT (n=6) animals. Scale bar: 50 μm. Results are shown as median (IQR); Kruskal–Wallis test followed by Dunn’s post hoc multiple comparisons test, ^*^p < 0.05, ^**^p < 0.01. **E:** Integrated density (arbitrary units or AI) of CD3^+^-lymphocyte infiltration quantified in naïve (n=4) and at 72 h after ischemic stroke in RjHan-NBT (n=2), RjHan-BT (n=6), RccHan-NBT (n=2), and RccHan-BT (n=6) animals. CD3 (green), RECA-1 (grey), and Topro-3 (blue). Scale bar: 50 μm. Results are shown as median (IQR); Kruskal–Wallis test followed by Dunn’s post hoc multiple comparisons test, ^*^p<0.05.

BT enhanced the systemic inflammatory response in both strains, but the effect was more pronounced in RjHan, suggesting that this sub-strain presents an earlier and more robust immune activation detectable at 72 h. Nevertheless, given that only a single time point was analyzed, it remains unclear whether RccHan develops a delayed response that peaks later or a generally attenuated one.

In the ischemic brain, granulocyte infiltration increased at 72 h in both strains as a consequence of stroke, but BT further augmented this infiltration only in RjHan rats. Regarding T cells, significant accumulation within the lesion was observed exclusively in BT groups of both strains, with a stronger effect in RjHan (Fig. 3D). These results suggest that BT acts as a driver of both peripheral and central inflammation, with RjHan rats being more sensitive to this effect (Fig. 3E).

Overall, our findings indicate that RjHan animals exhibit a more robust inflammatory phenotype at 72h after stroke, both in blood and within the lesion, particularly when bacterial translocation occurs. While this pattern is consistent with a faster inflammatory response in RjHan, confirmation of differential kinetics will require additional time points and functional markers.

## Discussion

This pilot study was designed as an initial step to investigate the effects of IS on IBP, BT, and their impact on peripheral and central inflammation, as well as stroke outcome. Stroke remains a leading cause of death and disability worldwide, and growing evidence indicates that systemic and immune mechanisms critically shape its pathophysiology and recovery [1–3]. Beyond the well-characterized central inflammatory cascades [2,3], stroke induces profound alterations in the immune system, including the phenomenon of CNS injury–induced immune deficiency [4]. More recently, IBP disruption, gut microbiota dysbiosis, and BT have emerged as central determinants of post-stroke immunity and outcomes [5–11,21].

In the present study, we compared Wistar rats obtained from two different suppliers, Janvier (RjHan) and Envigo (RccHan), and identified consistent differences in IBP, BT, systemic and cerebral inflammation, and stroke outcomes. Although both sub-strains are classified as Wistar, they have been maintained in separate breeding colonies for decades. Long-term isolation combined with vendor-specific husbandry practices likely contributed to genetic drift, epigenetic divergence, and distinct microbial exposures [18–20,23,24]. These accumulated differences may account for the variation observed in infarct size [15–17], immune activation [3,4], and bacterial dissemination [5,6], highlighting that even within a single nominal strain, vendor-related factors represent a significant source of experimental variability. While previous studies have examined strain-dependent differences in stroke outcomes [12–15], our data underscore that even sub-strains from different suppliers can respond divergently, emphasizing the importance of vendor selection for reproducibility and accurate interpretation of stroke-induced immune and gut– brain axis responses.

A particularly novel aspect of our findings is the demonstration of differential BT patterns between the sub-strains. As reported in the results, RjHan rats exhibited bacterial dissemination predominantly localized to the MLN, whereas in RccHan rats, bacteria were more broadly distributed across the spleen and liver. Analysis of the total bacterial burden outside the intestine showed no statistically significant differences between sub-strains; however, there was a clear trend toward higher overall dissemination in RjHan rats, which became statistically significant when examining bacterial loads specifically in the MLN. Additionally, taxonomic profiling revealed marked differences in the phyla translocated beyond the intestine: RjHan rats were dominated by *Bacteroidota*, followed by *Pseudomonadota*, while RccHan rats lacked *Bacteroidota*, with *Pseudomonadota* predominating and a pronounced reduction in *Bacillota* and *Actinomycetota* . Importantly, these differences persisted despite a 14-day acclimation period under standardized housing and diet, indicating that vendor-related microbiota characteristics remain distinct.

When combined with MRI data showing greater intestinal barrier permeability in RjHan rats [7], these findings suggest that the two sub-strains may preferentially use different routes of bacterial translocation. RjHan rats appear to rely primarily on a lymphatic route, leading to MLN colonization, whereas RccHan rats favor portal venous translocation toward the liver and spleen. This distinction aligns with the observed dissemination patterns and indicates that post-stroke gut-derived immune activation may proceed through different anatomical pathways depending on the sub-strain, likely influenced by vendor-specific background biology [8,9,18– 20].

Interestingly, the distinct patterns of bacterial dissemination observed between the two sub-strains, being more concentrated in the MLN of RjHan rats, suggest that this localized translocation triggers a stronger immune response. In contrast, in RccHan rats, bacterial dissemination is more diffuse but less dense per organ, potentially resulting in a qualitatively different inflammatory profile. Consistent with this, immune profiling revealed that RjHan rats mounted a more robust systemic and central inflammatory response at 72h post-stroke, particularly in the presence of BT. In peripheral blood, RjHan-BT animals showed pronounced shifts in lymphocyte and monocyte populations, with increased CD4^+^ and CD8^+^ T cells and decreased B cells and monocytes, whereas RccHan rats exhibited more modest or absent changes. In the ischemic brain, granulocyte and T cell infiltration was similarly enhanced in RjHan-BT rats, while RccHan rats displayed smaller increases. Collectively, these findings indicate that RjHan rats respond to stroke and BT with faster and more intense immune activation, whereas RccHan rats may exhibit a delayed or attenuated response, highlighting strain-dependent differences in both the kinetics and magnitude of post-stroke immune responses.

Beyond their mechanistic relevance, our observations underscore the importance of considering vendor origin in preclinical stroke research. Vendor-related differences in rodent behavior, microbiota composition, immune phenotypes, and pharmacokinetics have been reported [16–20], and our study extends these observations to include IBP, BT patterns, and stroke-related immune responses. This has important implications for reproducibility, as experimental outcomes obtained in one laboratory using Wistar rats from a single supplier may not fully extrapolate to animals from another source.

In conclusion, our study indicates that differences in stroke outcomes between RjHan and RccHan rats may reflect underlying genetic and microbiological divergence arising from separate breeding colonies [7,22–24]. These differences extend to intestinal barrier integrity, patterns of bacterial dissemination, and the resulting immune responses. Notably, RjHan rats exhibit more pronounced IBP disruption and lymphatic-driven BT, whereas RccHan rats show broader dissemination consistent with portal translocation [7–9]. These findings emphasize the need to carefully consider vendor origin when designing and interpreting studies of the gut–brain axis and post-stroke immunity, as such differences may critically influence both pathophysiological insights and translational validity.

This study has several limitations. First, analyses were restricted to a single time point (72h), precluding definitive conclusions regarding the kinetics of immune responses and bacterial translocation. Second, the sample size was modest, reflecting the exploratory nature of this pilot study. Third, functional markers of immune cell activation, trafficking, and proliferation were not assessed, limiting mechanistic interpretation. Finally, BT was inferred solely by microbial culture, without molecular confirmation. Addressing these limitations in future studies will be essential to validate and expand upon our findings.

## Acknowledgments

This research was funded by grants from the Spanish Ministry of Science and Innovation (PID2020-117765RB-I00; Dr Pradillo; PID2022-140616OB-I00, Dr Moro), Leducq Trans-Atlantic Network of Excellence (TNE-21CVD04; Drs Lo, Moro, Lizasoain), Instituto de Salud Carlos III, the European Development Regional Fund, RICORS-ICTUS (RD21/0006/0001), and the FORTALECE program (FORT23/00023; Dr Lizasoain).

