## Supplemental Material for "Sub-strain-Dependent Differences in Gut Barrier Permeability, Bacterial Translocation and Post-Stroke Inflammation in Wistar Rats"

**ONLINE SUPPLEMETAL MATERIAL**

**METHODS**

**Experimental groups, focal cerebral ischemia and neurological evaluation**
All experiments were performed in a randomized (coin toss) and blinded manner. The experimental groups consisted of naïve (n=4) rats and RjHan and RccHan Wistar rats subjected to middle cerebral artery occlusion (MCAO, n=8 per experimental group).
All the surgeries were conducted under anesthesia with isoflurane in a mix of O_2_ and N_2_O (0.2/0.8 L/min) and body temperature was maintained at 37.0±0.5°C using a servo-controlled rectal probe–heating pad. In rats, a tandem permanent occlusion of the left common-carotid artery by ligature and the left MCAO by electrocoagulation was performed as previously described [1]. After analyzing bacterial growth in organs, all groups were further classified as having bacterial translocation (BT) or not (NBT).

The neurological function was evaluated in naïve animals, and at 72h after surgery in the MCAO groups by motor and behavioral scales as described previously [2].
First, for the motor evaluation, a motor scale described by Hunter and colleagues in 2000 was used [3]. According to this scale, the animal receives a score of 0 points if no motor deficits are present. A lack of full extension of the right forelimb corresponds to 1 point; a decreased grip in that limb while the animal is held by the tail is scored as 2 points; circular movement on the forelimbs toward the contralateral side when held by the tail indicates 3 points; walking in circles toward the contralateral side when stimulated corresponds to 4 points, and the absence of response to stimuli is scored as 5 points. On this scale, a higher score indicates a greater motor and neurological deficit.

On the other hand, a behavioral scale described by Madrigal et al. in 2003 was used to assess behavioral/motor status by scoring various parameters [4]. In this scale, animals with greater behavioral deficits receive lower scores. The evaluated aspects were: body posture (0–4 points), spontaneous activity (0–3 points), arousal response when transferred to a new cage (0–5 points), crawling (0–3 points), escape response when touched (0–3 points), and the ability and strength of grip with the forelimbs (0–4 points).

**Experimental infarct size and GBD determination by MRI**

Infarct volume was assessed in all MCAO animals 72h after experimental IS using T2-weighted MRI (Icon 1T-MRI; Bruker BioSpin GmbH, Germany). After anesthetizing the animals as previously described, T2-weighted 3D images were acquired with the RARE technique (TR = 2200 ms; echo train length = 12; echo spacing = 15 ms; TE = 75 ms; averages = 1; FOV = 34 × 31 × 12 mm; matrix = 136 × 124 × 34; resolution = 0.25 × 0.25 × 0.50 mm) in a total time of approximately 10 minutes. Infarct size was expressed as a percentage of the ischemic hemisphere (%IH) as previously described [5].

Intestinal barrier permeability was assessed in naïve rats and untreated rats 72h after IS using T1-weighted abdominal MRI with orally administered contrast agents. Imaging was performed on a Bruker Biospec 4.7T MRI system (Bruker, Ettlingen, Germany) with a 12 cm gradient and a 7 cm RF coil. To optimize image quality, animals fasted for 12 hours and were anesthetized during scanning, with temperature and respiration continuously monitored. One hour before imaging, rats received by oral gavage 1.5 ml of mannitol (Manitol Mein 10%, Fresenius Kabi S.A.U) and 0.5 ml water as osmotic agent, along with 250 mg/kg of MnCl₂·4H₂O (CAS 13446-34-9, PanReac, Barcelona, Spain) dissolved in 1.5 ml of water as contrast agent. Respiration-synchronized 2D spin-echo T1-weighted images with fat suppression were acquired (TR = 802 ms; TE = 10 ms; 8 averages; FOV = 12 × 6 cm; matrix = 256 × 128; slice thickness = 1 mm; 12 slices; resolution = 469 µm²; scan time ~14 min). Finally, the acquired images were analyzed using ImageJ 1.41 software (NIH, Bethesda, MD, USA) to quantify intestinal barrier damage by detecting contrast agent leakage into surrounding tissues.

**Analysis of BT by microbiological cultures**

To investigate BT after stroke, microbiological cultures of mesenteric lymph nodes (MLN), liver, spleen, and lungs were performed in naïve and MCAO animals 72h post-surgery. Under terminal anesthesia, samples were aseptically collected, weighed, and homogenized in sterile saline. Homogenates (undiluted and diluted) were plated on Columbia, MacConkey, and Brucella agar (all from Becton, Dickinson and Company, NJ, USA). Columbia and MacConkey plates were incubated at 35°C for 48h under aerobic conditions for the growth of strict aerobes. Brucella agar plates were incubated at 35°C for 96h in an anaerobic chamber to support the growth of strict and facultative anaerobic bacteria. After incubation, colony-forming units (CFUs) were counted and identified by stereomicroscopy, culture, and API biochemical tests. Images were taken with a SteREO Discovery V.8 stereomicroscope and processed with Zen software. Distinct colonies were marked, subcultured, and further identified. Results were reported as log₁₀ CFU/g tissue (detection limit: 1.3 log₁₀ CFU/g). Bacterial phyla were classified according to the International Code of Nomenclature of Prokaryotes (ICNP) [6].

| **New nomenclature** | **Old nomenclature** |
| --- | --- |
| *Bacillota* | *Firmicutes* |
| *Bacteriodota* | *Bacteroidetes* |
| *Actinomycetota* | *Actinobacteria* |
| *Pseudomonadota* | *Proteobacteria* |
| *Fusobacteriota* | *Fusobacteria* |

**Determination of peripheral and central inflammation**

Peripheral inflammation was evaluated by flow cytometry in *naïve* animals and in MCAO animals at 72 h post-surgery, using samples from bone marrow (BM) and blood. The populations analyzed included CD4⁺ and CD8⁺ T cells, B cells, monocytes and granulocytes. Following sample collection, BM samples were mechanically homogenized in 1% PBS-BSA and filtered through a 50 µm mesh. For both BM and blood samples, red blood cells were lysed, Fc receptors were blocked, and cells were centrifuged. Samples were then incubated with the appropriate antibody cocktails (detailed below), analyzed on a FACSCalibur cytometer (200,000 events per sample), and processed using FlowJo software. Data were expressed as percentages of total cells.

| **Antibodies for Rat Flow Cytometry** | | | |
| --- | --- | --- | --- |
| Cell population | Antibody | Fluorochrome | Supplier (Catalog No.) |
| CD4 T cells | CD45 | FITC | BioLegend (Cat. No. 202205) |
|  | CD4 | PE | BioLegend (Cat. No. 550002) |
|  | CD3 | PE | BioLegend (Cat. No. 201507) |
| CD8 T cells | CD45 | FITC | BioLegend (Cat. No. 202205) |
|  | CD3 | PE | BioLegend (Cat. No. 201507) |
|  | CD8a | APC | BioLegend (Cat. No. 201705) |
| B cells | CD45 | FITC | BioLegend (Cat. No. 202205) |
|  | CD24 | PE | BD Biosciences (Cat. No. 562104) |
| Monocytes | CD45 | FITC | BioLegend (Cat. No. 202205) |
|  | CD11b/c | APC | BioLegend (Cat. No. 201809) |
| Granulocytes | Anti-rat granulocytes | PE | BD Biosciences (Cat. No. 554905) |
|  | MHC Class II (I-Ek) | APC | Miltenyi Biotec (Cat. No. 130-107-874) |

To assess central inflammation, *naïve* and MCAO animals (72 h post-ischemia) were transcardially perfused, and brains were collected, fixed, sectioned (30 µm), and processed for immunofluorescence. Free-floating sections were stained with antibodies against granulocytes, CD3⁺ T cells, and the endothelial marker RECA-1, followed by Alexa Fluor 488- or Cy3-conjugated secondary antibodies. Nuclear counterstaining was performed with TO-PRO-3. For each animal, four random images were acquired from five consecutive brain sections (spaced 720 µm, covering +1.70 to −0.40 mm relative to bregma). Images were collected as Z-stacks at 20× magnification using a Zeiss LSM710 confocal microscope and analyzed by densitometry with ImageJ software. Granulocyte infiltration was quantified as the number of cells per mm². For infiltrating CD3⁺ T cells, the integrated density was determined as the ratio between the intensity of positive cells and the analyzed area.

**Online Suplemental References**
